# Automated acoustic classifiers provide insights into calling patterns of cicadas in tropical Australia

**DOI:** 10.64898/2026.09.17.752521

**Authors:** Yucheng (Scarlett) Zhu, Lin Schwarzkopf, Myles H. M. Menz, Slade Allen-Ankins

## Abstract

Insects are a key group in ecosystems and economics, but are increasingly under threat, making ongoing monitoring essential to preserving them. Traditional monitoring approaches are time consuming, among other limitations. Automated acoustic recognition can be a good solution for monitoring arboreal, soniferous insects, such as cicadas, and the acoustic biology of cicadas in Australia is understudied, especially using automated methods. To evaluate the applicability of automated approaches for detecting cicadas, we used the deep-learning acoustic model BirdNET (v2.4) to perform embedding searches for *Cystosoma schmeltzi* (lesser bladder), *Illyria burkei* (eastern rattler), *Macrotristria intersecta* (corroboree cicada), *Thopha sessiliba* (northern double-drummer), and *Pauropsalta opaca* (fairy dust squawker), using audio recorded by Passive Acoustic Monitoring over 15-months at 18 sites in north Queensland. All classifiers achieved high area under the precision-recall curve (AUPRC) ranging from 0.753 to 0.982. We used a precision-prioritised threshold (≥ 0.85) to constrain the false positive rate while retaining adequate outputs, which were consistent with the documented species ecology and life history. This approach was readily applicable to the Cicadidae, offering a scalable solution for biodiversity monitoring in complex acoustic environments.

## Introduction

Insects play key ecological and economic roles (Lavelle et al. 2006; Landis et al. 2008; Schowalter 2016), and a diverse community is essential for ecosystem sustainability (Worm and Duffy 2003; Stachowicz et al. 2007; Traill et al. 2010). However, insects are increasingly under threat and declining world-wide (Wagner 2020). Accordingly, monitoring for insects is crucial for biodiversity assessment and quantifying ecosystem function, and to help understand threats to these vital groups.

Despite its importance, common monitoring methods for many insect taxa (e.g., pit trapping, malaise trapping, and hand capture with manual identification) can present challenges for arboreal insects (Mori et al. 2026), and thus we require an effective tool to detect the biodiversity of this assemblage. The use of acoustic sensors to conduct environmental monitoring (i.e., Passive Acoustic Monitoring or PAM) holds great promise for advancing the surveillance and monitoring of cryptic animals that call. Species-specific vocalisations have been described for many vocal insects (Mankin et al. 2011), and substantial research has been conducted on insect acoustic signalling, yet automated detection remains underrepresented. This is partly because calling of soniferous taxa is species-specific, but there is a scarcity of curated reference recordings with reliable metadata. Despite the challenges, insect calls may offer reliable signals indicating their occurrence and activity. For example, cicadas are vocal insects that call loudly during their mating season, which is typically strongly seasonal (Wolda 1989). This calling behaviour is tightly linked to habitat use and the timing of emergence, and therefore provides a measurable proxy for their presence (Young 1972; Rodenhouse et al. 1997; Sueur 2002; Takakura and Yamazaki 2007). From an ecological perspective, detection at species level for cicadas offers a robust basis for practical investigations of biodiversity. However, traditional detection approaches, including manual identification of acoustic events, is time consuming (Digby et al. 2013). By contrast, automated classifiers offer a scalable solution for detecting soniferous animals in extensive recordings over long term studies.

The deep learning system BirdNET (Kahl et al. 2021), has been widely applied to birds (Doohan et al. 2026), and several studies have demonstrated that it performs well detecting distinctive signals for other taxonomic groups (e.g., whales (Tolkova et al. 2025), amphibians (Leung et al. 2026), and reptiles (Bartholomew et al. 2025)). All these species emit distinctive single calls, but to use this system for insects, it is critical to determine whether BirdNET performs well at detecting continuous acoustic streams that are minimally modulated and spectrally monotonic, such as those emitted by most vocal insects, perhaps most famously by cicadas (Hemiptera, Cicadoidea).

We assembled acoustic data collected as part of another study, using passive acoustic monitoring (PAM) conducted in blocks of approximately 10 days, over 15 months. We developed five, species-specific cicada recognisers using the deep-learning acoustic model BirdNET to perform embedding searches for target cicada species, followed by iterative model training using manual validation at the clip level. We then summarised the automated detection outputs describing species’ diel and seasonal calling activities. This study aimed to: 1) Develop effective automated classifiers capable of processing long duration ecoacoustic datasets, to detect vocalisations of the target cicada species. 2) Evaluate classifier performance to explore the feasibility and potential of automated recognition in insect bioacoustics. 3) Investigate the cicadas’ seasonality, distribution across the study sites, and diel calling patterns during our survey periods using the automated detection output.

## Methods

### Acoustic Data collection

We used existing audio data collected from a Gulf Savannah Natural Resource Management (NGM) Biodiversity Project, which was originally designed to detect effects on vertebrate biodiversity of grazing. Eighteen sites were surveyed: six paired sites on cattle grazing stations (two each on Dagworth, Eveleigh, Mount Surprise, Ooralat, Van Lee and Whitewater Cattle Stations), and six sites in two Indigenous Protected Areas (IPAs, three each on Undara and Talaroo) in the tropical Gulf Savanna of northern Queensland, Australia (Figure 1). At each site, one Long Term Bioacoustic Recorder (BAR-LT™, Frontier Labs) was secured on a tree ∼1.5 m above the ground and set to record 1-hr files 24 hours a day in 16-bit depth at a sampling rate of 22.1 kHz in .wav format. Recording was conducted over four survey periods, including one Austral summer and winter season, and two spring seasons from September to October in 2022 and 2023 respectively (Table 1), yielding approximately 24,000 hours of recording.

**Table 1.** The time and site deployment for each of the four survey trips. Details are based on Brodie et al. (2023).

| Deployment | Recording period | Recording sites |
| --- | --- | --- |
| July, August 2022 | 18/07/22 - 31/07/22<br>(14 days in the dry season) | Dagworth (2 sites),<br>Whitewater (2 sites),<br>Van Lee (2 sites),<br>IPA Talaroo (3 sites). |
|  | 03/08/22 - 18/08/22<br>(16 days in the dry season) | Eveleigh (2 sites),<br>Mt Surprise (2 sites),<br>Ooralat (2 sites),<br>IPA Undara (3 sites). |
| December 2022, January 2023 | 04/12/2022 - 07/01/2023<br>(6.8 - 15.9 days in the early wet season) | All sites except Ooralat and Talaroo (1 site). |
| July, August 2023 | 3/07/2023 - 8/08/2023<br>(9.5 - 18.7 days in the dry season) | All sites except one site in Talaroo. |
| September to November 2023 | 25/09/2023 - 02/11/2023<br>(16 - 18 days in the transitional season) | All 18 sites. |

**Figure 1.**
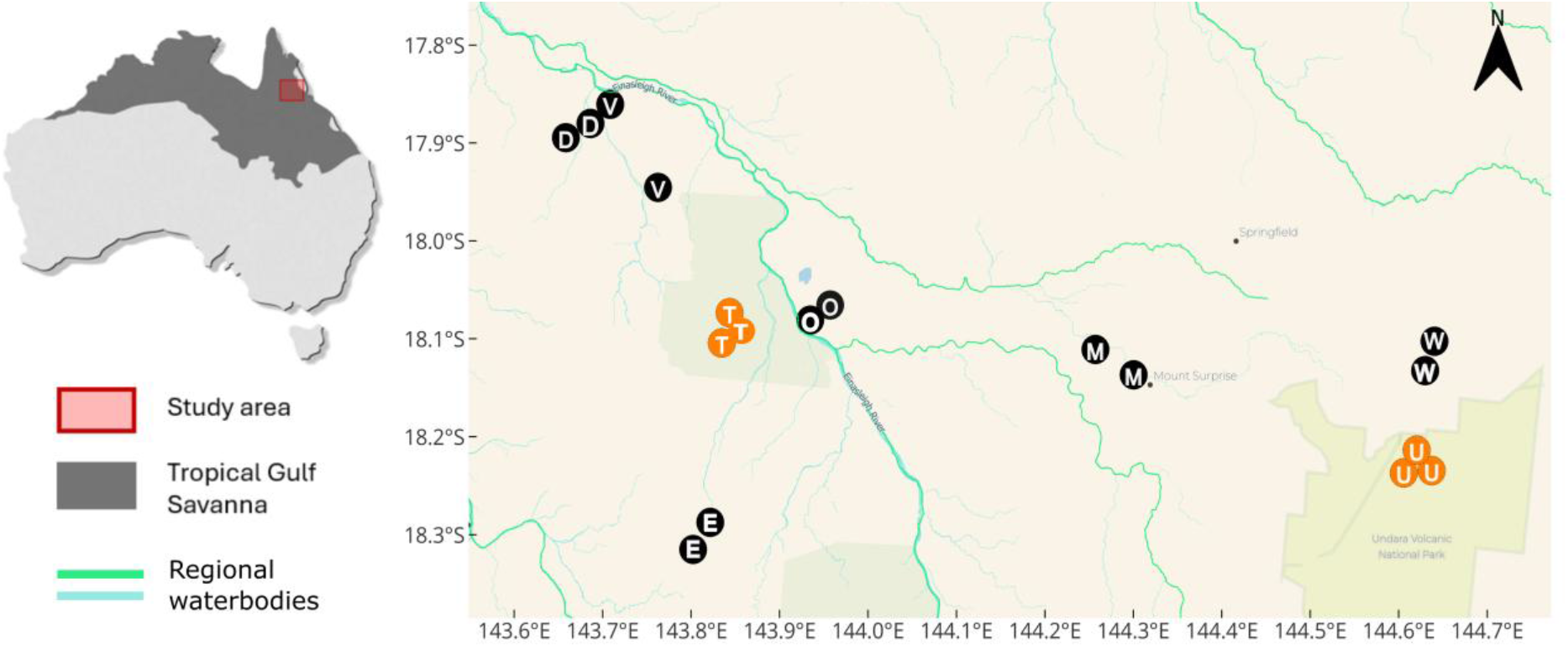
Study sites in the tropical gulf savanna, Northern Queensland, Australia. Black circles represent sites located on grazed properties, with two stations at each site. The letters on the circles are the initials of the property names: Dagworth (D), Eveleigh (E), Mount Surprise (M), Ooralat (O), Van Lee (V), and Whitewater (W) Cattle Stations. Orange circles stand for the sites in Indigenous Protected Areas (IPAs): Talaroo (T) and Undara (U). Green and blue lines indicate regional watercourses, including the Einasleigh River.

### Acoustic classifier construction

To develop acoustic classifiers for the automated detection of cicada calling, we used a four-stage methodological process, including: 1) obtaining example calls for each target species; 2) conducting a search of the embeddings from BirdNET for each location and time combination, using the example call; 3) manually validating the similar sounds uncovered by the search for the presence or absence of the target species; and 4) training linear classifiers using the labelled embeddings and evaluating performance on held-out data.

### Obtaining song examples

To select species of cicadas likely to inhabit the Gulf Savannah region, we filtered data in the Atlas of Living Australia platform (Belbin et al. 2021), using an area that included all 18 study sites, plus a 10 km buffer. We then collated a list of cicada species that had been observed inside this geographic area, where the five target species were selected as the most common species in the community, also their vocal presence was identified by manual verification and expert validation. Example calls for the target cicadas were downloaded from Popple (2026), a web-based repository of Australian cicada sounds. The website provides professionally verified recordings of each Australian cicada species.

The song examples sourced from Popple (2026) were excluded from the final classifier training set.

### Embeddings search

Example audio, and unlabelled recordings collected from our study area, were divided into 3-second segments, for selecting corresponding feature embeddings, which provide a multivariate representation of an audio segment (Stowell 2022). The search was used to provide examples that could later be used to build classification models that allow automated call detection in acoustic events (McGinn et al. 2023; Allen-Ankins et al. 2025).

The feature embeddings of both the exemplar calls of cicadas and the unlabelled audio dataset to be searched were processed with the *embeddings*.*py* script from the BirdNET Analyzer v2.4 (Kahl et al. 2021), which produced a 1024 length embedding vector representing the acoustic structure of each 3-second window; for an audio example this was a single vector, for the audio being searched it was vectors for as many 3-second segments as were contained in the sample recordings (Allen-Ankins et al. 2025) (typically about 3-500,000 clips in our data).

To determine the similarity between the embedding vector of the target cicada vocalisation and each embedding vector in our unlabelled recordings, we calculated the Euclidean distance between the two vectors using R version 4.1.2 (R Core Team 2021). This produced a distance value for every 3 seconds of unlabelled audio, such that more similar embeddings had lower Euclidean distance values (Allen-Ankins et al. 2025).

Once these distances had been calculated, approximately 150 3-second segments were selected from across the range of distances, for manual labelling. This process ensured that the manually reviewed segments spanned the full continuum from highly dissimilar to highly similar acoustic samples. The selected segments were validated and annotated as either a true positive (containing a target call) or a false positive (nontarget sounds) using Audacity software (Washnik et al. 2023), which shows spectrograms, in addition to allowing listening to the potential target signals. This audio-visual validation was helpful for comparison and confirmation of whether target signals did occur within soundscapes. In addition, we provided examples of calls from our data to an expert (Dr. Lindsay W. Popple), to further validate our species identifications.

### Model training and performance evaluation

A classifier was implemented as a binomial generalised linear model trained using the 1024 length embeddings vectors as predictors, and call presence or absence as the response. The validation results from the embeddings search were used to train initial models for each species. The training process required sufficient training data to allow high performance to be achieved in automated acoustic classification. Acoustic classifier performance is influenced by the distinctiveness of the target vocalisation relative to co-occurring sounds, as well as by the amount and quality of the training data supplied (Wood and Kahl 2024). Several training cycles were required for each species, and the performance of each classifier was manually evaluated to determine whether further improvement was warranted.

To evaluate classifier performance, both precision and recall were quantified across a range of threshold scores, generating a precision–recall curve. Model precision is defined as the proportion of classified detections that truly contain a desired vocalisation (1 – false positive rate), whereas model recall is the proportion of all vocalisations present in a recording that are successfully detected by a classifier (1 – false negative rate). The area under the precision-recall curve (AUPRC; scored from 0 to 1) provides a useful summary metric for classifier evaluation (Knight et al. 2017), specifically measuring the trade-off between precision and recall. High-performing classifiers maintain both high precision and high recall, producing greater AUPRC values. For each classifier, AUPRC was calculated using the *yardstick* package (Saito and Rehmsmeier 2017) in R v4.5.0 (R Core Team 2025), based on performance evaluated on approximately 500 audio segments. Classifiers were iteratively improved by incorporating additional training recordings until an AUPRC value greater than 0.75 was achieved.

A classifier was considered to have acceptable performance if it satisfied the following criteria: a high AUPRC (>0.75), a clearly defined threshold logit score that effectively separated true positives from false positives, a reasonable recall rate (>0.5), and a high precision value (>0.85). Precision was prioritised because cicadas are chorusing species, and it is not important to detect every 3-second segment including a call, as long as a reasonable number are detected. Thus, false positive detections would have a greater impact on the validity of the results than false negatives. This approach ensured that false positive rates remained low, while false negative rates were kept approximately consistent among recordings. Once a classifier was deemed acceptable, a score threshold was selected, above which audio segments were classified as detections, fully automating the acoustic detection process, although we did haphazardly check some calls detected at unusual times, when few or no other calls were detected, these were all confirmed to be actual calls.

To estimate diel calling patterns, the call detections were aggregated by hour and analysed for each species using binomial hierarchical generalised additive models (HGAMs) in R using the *mgcv* package (Wood 2011). The proportion of detected audio segments within each hour was modelled using a binomial distribution with a logit link. Diel variation in calling activity was represented by a cyclic smooth of hour that was allowed to vary for each site and the model contained a random effect to account for day-to-day variation due to repeated observations within sites. Predictions from the fitted HGAMs were used to plot site-specific and overall diel calling patterns for each species.

## Results

### Details of calls of individual cicada species

Using the methods described above, we described the calls of five cicada species at the study sites.

Corroboree cicadas (*Macrotristria intersecta*) call in a harsh, fizzing whine with a continuous tonal component that audibly switches between brighter and duller, occupying broad frequency spectra ranging from 3.3 to 11 kHz (Figure 2a, b).

**Figure 2.**
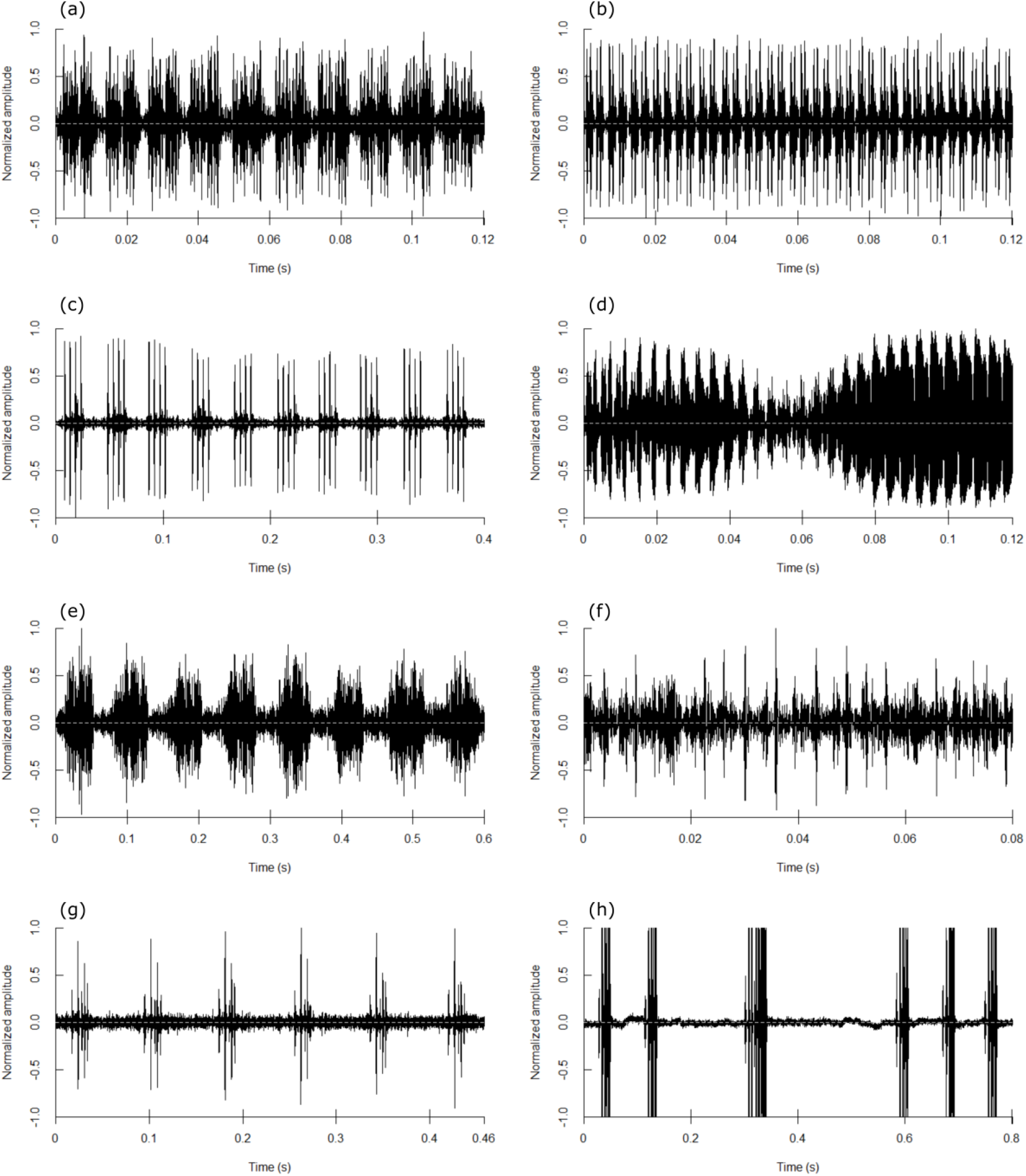
Oscillograms of the songs of five cicada species. Note the time period shown on the Y-axis varies among species to allow inspection of the pulse rate in each case. (a, b) *Macrotristria intersecta* alternates between two call phases shown by different sound patterns in its repertoire. (c) *Illyria burkei*, (d) *Thopha sessiliba*, (e) *Cystosoma schmeltzi*, and (f-h) *Pauropsalta opaca*, with its song components 1,2 and 3.

Vocalizations of the eastern rattler (*Illyria burkei*) exhibited simpler acoustic characteristics in soundscapes. Their song was a continuous, and relatively high-pitched, rattling buzz with frequencies ranging from 4.2 to 11 kHz (Figure 2c).

Northern double-drummers (*Thopha sessiliba*) produce a loud, electric whine, fluctuating regularly between strong and secondary accents across changing pitches in a continuous buzz. The frequency covered a broad range with a dominant band between 4.2-6.5 kHz (Figure 2d).

Lesser Bladder cicadas (*Cystosoma schmeltzi*) vocalized as a broken rattle in a regular chanting call, or as a guttural growl. Their song was not completely continuous and contained brief pauses. The sound frequency spanned from 2.2 -10.5 kHz (Figure 2e).

The song of Fairy Dust Squawkers (*Pauropsalta opaca*) consisted of three components. The first component was a long buzz. The second component contained a series of chirps, emitted rapidly, with each short series connected by a single click. The third component was a longer, continuous trill that could last for several seconds. Sometimes the longer buzz was produced between a successive series of rapid chirps. The song possesses a broad frequency band (1-11 kHz) (Figure 2f, g, h).

### Classifier performance

Across these five species, the classifiers achieved high overall performance (AUPRC median = 0.856, Table 2). Using a precision-prioritised threshold (≥0.85), species specific recall varied from 0.58 to 1 (Table 2). The precision-recall curves for each species’ model further illustrate performance differences and operating trade-offs (Figure 3).

**Table 2.**
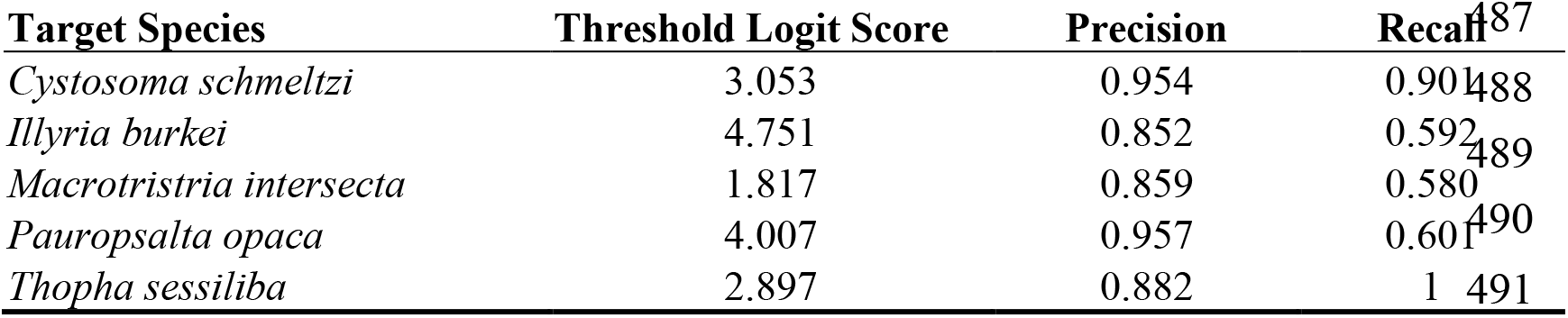
Performance metrics of species-specific automated classifiers. Threshold logit score corresponds to the precision prioritised threshold (precision ≥0.85).

**Figure 3.**
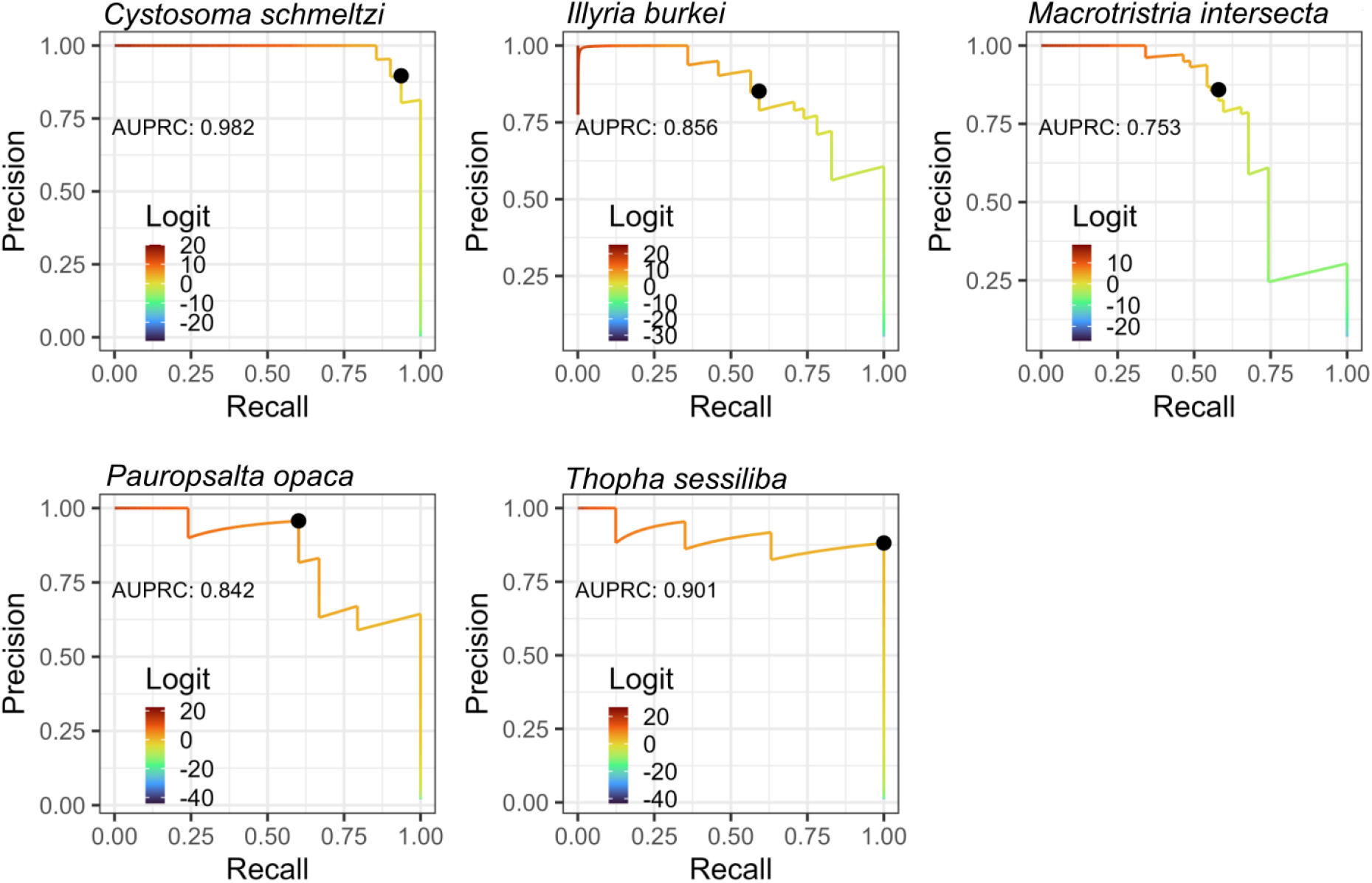
Species specific precision–recall curves and AUPRC values for calls of five cicada species. Each panel shows one target species. Lines depict precision as a function of recall across decision thresholds. AUPRC is annotated in each panel. The colour scale encodes the classifier logit score at each operating point. The black dots represent selected decision thresholds on the PR curves.

### Detection outputs of classifiers

Detections generated from the five classifiers were plotted, producing a diel calling pattern for each of the five cicada species (Figure 4a-e). This allowed us to summarize the distribution of seasonality (Table 3), across the study sites (Figure 5), and species-specific features of acoustic behaviours.

**Table 3.**
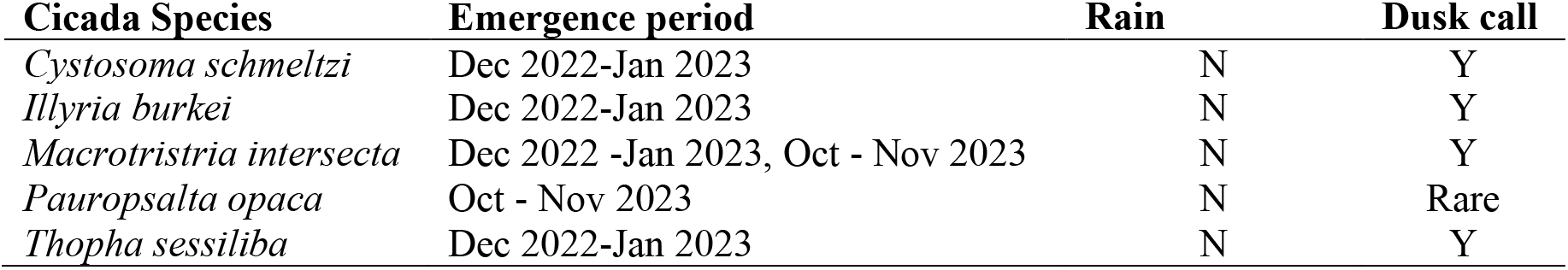
Detected seasonality, habitat distribution, and diel call activity of the five cicada species.

**Figure 4.**
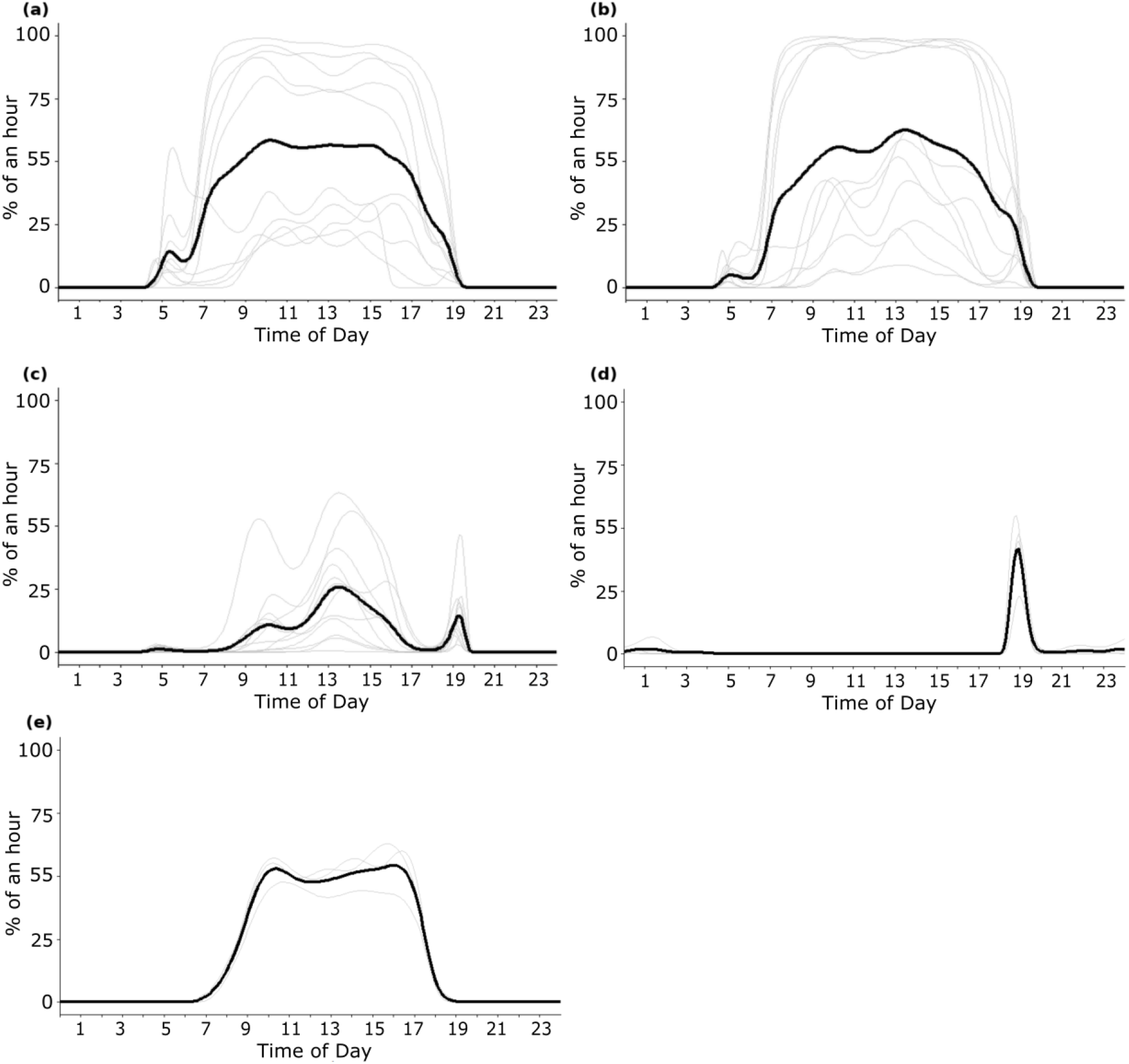
The calling rhythms of the cicadas across 24 hours during December 2022-January 2023, except for (e). Thin lines represent calling at individual sites, while the thicker black line is the median calling pattern among sites. (a) *Illyria burkei*, (b) *Macrotristria intersecta*, (c) *Thopha sessiliba*, (d) *Cystosoma schmeltzi*, (e) *Pauropsalta opaca*, the calling songs in the pattern were detected during October - November 2023.

**Figure 5.**
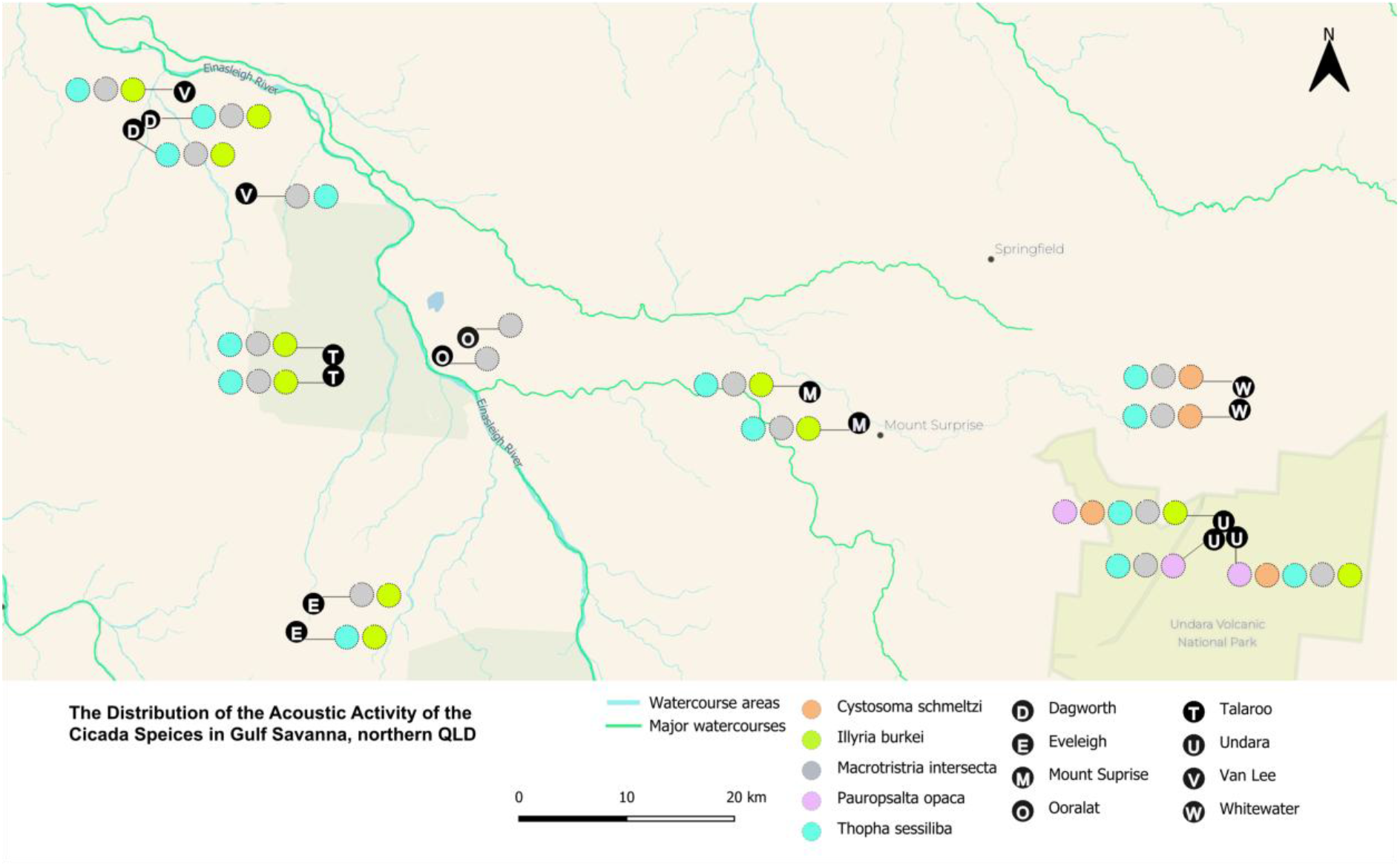
Distribution of the detected calls of the five cicada species. Dot colours stand for different species; black dots represent study sites, with the initial of the site name inside each dot.

### Diel calling patterns

Diel calling behaviour of *I. burkei* was strongly diurnal across sites (Figure 4a). Calling included a short prelude, generally commencing in the early morning, then increased rapidly and reached an intensive chorus around mid-morning, forming a pronounced mid-day plateau. Calling activity then gradually decreased in the afternoon and ceased around dusk.

Similarly, *M. intersecta* started calling very early, and then chorusing was typically continuous all day (Figure 4b). Notably, at some sites, both *M. intersecta* and *I. burkei* showed a break in calling after dawn. *M. intersecta* also ceased calling at some sites in the evening, before resuming around sunset. The amount of calling varied among sites in both species.

Calling by *T. sessiliba* was bimodal, starting in the early morning, followed by a period of silence. Chorusing then increased again after 08:00, reaching its highest intensity during mid-morning, sometimes declining in the late morning and then increasing again in the early afternoon (Figure 4c). They ceased calling in the late afternoon, then called again, stopping at in the early evening.

Compared to the other species, which chorus much of the day, lesser bladders (*C. schmeltzi)* typically called only in the evening (Figure 4d). Additionally, low levels of nocturnal calling were detected at some locations (Figure 4d).

Vocal activity of *P. opaca* was slightly bimodal diurnally across our sample days and sites (Figure 4e). Calling started in the morning, followed by nearly continuous activity until late afternoon. This species normally stopped vocalising in the evening.

## Discussion

Our study demonstrates that automated acoustic classifiers, created using an embeddings search, can reliably detect multiple co-occurring cicada species. Classifier performance was consistently high across all five taxa, enabling robust extraction of species’ calling behaviour from audio datasets. Each species exhibited distinct daily calling patterns. These findings illustrate that acoustic monitoring using an automated classifier can describe calling in cicadas at a fine scale, and provide a vivid picture of their vocalising patterns in tropical savanna landscapes.

## Development of the classifiers

We successfully developed recognisers for five Australian cicada species, direct from environmental recordings, using an embeddings search followed by transfer learning from the BirdNET model v2.4. This method has been used to detect vertebrates, with a high precision achieved (Enari et al. 2019; Leung et al. 2025; Allen-Ankins et al. 2026). However, it was not clear if this technique would work for insect choruses, which are continuous and minimally modulated compared to birds. In our study, this method identified species-specific signals with precision above 85% in empirical datasets for each focal species, even in the presence of overlapping sounds from other species, and environmental noise. Compared to the development of specialised convolutional neural networks (CNNs) (Okamoto and Oguma 2025), our models could be developed with comparatively minimal training effort, progressing from very few example calls to a well-performing trained model in a few validation iterations. In some locations in our dataset, similar frequency ranges were shared among different species in the morning, afternoon, and at dusk. Interspecific overlapping frequencies were properly discriminated *via* our classifiers, and signals emitted by species not of interest could be successfully eliminated.

## Spatiotemporal patterns of species’ vocal activity

Adults of Australian cicadas typically emerge during a specific season each year and are present only for a 2-to-6-month activity window (Popple 2026). The five studied cicada species all showed strong concordance with documented records regarding seasonality, diel calling patterns, and habitat preferences.

### Illyria burkei

In our data, *I. burkei* occurred in open forest and woodland on poor soils associated with box and ironbark eucalypts, similar to habitats documented by (Moulds 1985). The seasonal timing of calling in December and January was also consistent with their reported emergence period from November to January (Moulds 1985; Popple 2026). We found that populations vocalized throughout the day with a resurgence at dusk, consistent with patterns reported by Popple (2026).

### Macrotristria intersecta

In our study, *M. intersecta* inhabited open eucalypt woodlands, consistent with the previous habitat records (Popple 2026). We detected calling by *M. intersecta* from mid-October to early January, matching their documented emergence timing from September to February (Popple 2026). In our study, *M. intersecta* species called during the day and at dusk, as reported by (Ewart 2005).

### Thopha sessiliba

We detected *T. sessiliba* calling up to 1.1 km from tributaries of the Einasleigh River, Parallel Creek, and Cattle Creek systems, consistent with reports of *T. sessiliba* as restricted to watercourses (Popple 2026).

Adults of *Thopha sessiliba* called in summer from December to early January across most of our sites, which aligns with the species description as a typical summer species (Ewart 2005).

The diurnal calling cycle of the species ended with short peak at dusk. This was mirrored to the calling pattern reported by Ewart (2005) and Popple (2026).

### Cystosoma schmeltzi

In our study, *C. schmeltzi* vocalized in moist to dry open *Eucalyptus* forests and woodlands, consistent with previous records (Popple 2026), but was also detected in *Melaleuca* swampland, potentially extending its habitat associations. Vocal activity occurred during the Austral summer, as previously reported, but differed between years: the species was detected in 2022–23 but not during comparable sampling in the following season. This interannual variation may reflect differences in local weather or patterns of adult emergence (Coombs 1996; Kaliang et al. 2025).

Calling was concentrated around dusk, consistent with previous observations (Popple 2026), but sometimes occurred after dark, suggesting that *C. schmeltzi* is not strictly crepuscular (Young 1980, 1981b; Gogala and Riede 1995).

### Pauropsalta opaca

At our study site, *P. opaca* was found at the edge of its range and occurred only at our Undara sites. It called in eucalypt woodland in Austral summer and spring, and called diurnally, stopping before dusk, consistent with other records for the species (Popple 2026).

## Call overlap and temporal segregation

Our results showed substantial temporal overlap in calling activity among sympatric cicada species, suggesting that diel partitioning is not the primary mechanism of acoustic niche separation. An exception occurred during the avian dawn chorus. Both manual inspection and recogniser detections indicated that *M. intersecta* reduced or paused signalling during this period, when bird songs overlapped its dominant frequency range (∼5–8 kHz). This may reduce acoustic masking (Brumm and Slabbekoorn 2005; Anjana and Teji 2025). In contrast, *I. burkei*, whose signals are centred at the higher frequency range of 7–11 kHz, showed less interruption during the dawn chorus. This pattern suggests that cicadas may avoid calling due to frequency-dependent interference from birds, just as birds have been documented to avoid calling with cicadas when their dominant frequencies overlap substantially (Berman et al. 2024).

During the day, *M. intersecta* and *I. burkei* frequently chorused simultaneously where they occurred sympatrically. This contrasts with expectations that co-occurring species reduce acoustic interference through temporal partitioning (Leroy 1977, 1978; Riede 1997; Sueur 2002). Their dominant frequencies, however, are well separated, likely allowing simultaneous signalling with little interference. Temporal segregation is probably unnecessary when sympatric species occupy distinct frequency bands (Ewart 2001). Moreover, shared ecological constraints, including microclimate and predation risk, may restrict calling to similar favourable periods, encouraging temporal overlap (Henwood and Fabrick 1979; Sanborn et al. 1995).

Temporal overlap also occurred among *M. intersecta, I. burkei*, and *T. sessiliba*, even though the broad frequency range of *T. sessiliba* (3.3–11 kHz) overlapped both other species. Where frequency and temporal partitioning are limited, interference may instead be reduced through differences in signalling behaviour, including spatial segregation or differences in pulse structure (Sueur 2002).

At dusk, these three species and *C. schmeltzi* called concurrently, producing a particularly complex acoustic environment. Concentrated dusk calling may reflect favourable conditions for sound transmission (Van Staaden and Römer 1997), as well as ecological factors such as acoustic crypsis and avoidance of predation risk (Sanborn et al. 1995; Oliveira et al. 2021). Effective communication under these conditions is likely facilitated by fine auditory discrimination among species (Fonseca et al. 2000). Occasional nocturnal calling by *C. schmeltzi* may further reduce both predation risk and interference from diurnal soniferous insects (Young 1981b; Sueur 2002; Tomita 2021).

Overall, our results suggest that these sympatric cicadas rely less on temporal partitioning than on differences in signal frequency and signalling behaviour to maintain effective intraspecific communication. Such multidimensional acoustic niche separation may allow species to chorus simultaneously, even when some overlap in signal frequency occurs.

## Breaks in calling

In our study, *T. sessiliba* exhibited distinct silent periods in its calling activity, with fewer vocalizations for approximately two hours after the dawn chorus began, and another two hours during the dusk chorus. Similar periods of reduced or absent calling were also observed at some sites for *M. intersecta* and *I. burkei*. Comparable silent periods have been reported for two Australian cicada species (Ewart 2001) and nine sympatric cicada species in a Mexican rainforest (Sueur 2002), suggesting that this pattern is widespread among cicadas. Reduced calling following the initiation of the dawn chorus may reduce exposure to predators, as it coincides with a time of high activity by insectivorous birds (Alexander 1960; Doolan and Mac Nally 1981; Young 1981), while the silent period associated with the dusk chorus may reflect a recovery period before an energetically demanding bout of evening calling (Sanborn et al. 1995).

## Vocalizations in precipitation

Our results show that rainfall failed to suppress calling activity. Although the literature reports that cicadas call in clear weather, and that they cease calling in rain (Fonseca and Allen Revez 2002; Hart et al. 2015), all our species were detected calling under light, moderate, and heavy rain conditions.

While conducting manual inspections, louder choruses were observed when the rainfall escalated to a heavier pour (Figure A8). This is important, because some studies have recommended suspending observations during rain (Hart et al. 2015; Strang 2025). From our observations, we suggest that avoiding observations of cicada calling during rainfall would be counterproductive for these species.

## Conclusion

The automated acoustic monitoring approach was highly effective at detecting cicada vocalisations within complex natural soundscapes. Embedding-based search methods further enabled the rapid identification and validation of calling events, demonstrating the capacity of passive acoustic monitoring to detect insect species reliably under real-world conditions. Simple linear classifiers trained on deep-learning embeddings successfully identified target insect species despite the presence of non-target biological sounds, anthropogenic noise, and weather-related signals. These results highlight the value of automated acoustic methods for studying insect behaviour at scales and levels of environmental complexity that would be difficult to achieve using traditional field observations alone.

## Acknowledgements

We thank Dr. Lindsay W. Popple for his assistance validating our cicada species identifications. We thank Dr. Sheryn Brodie for providing information on recorder deployment throughout the survey periods. We also thank Gulf Savannah NRM for providing funding for the project (N. Gobius) and conducting fieldwork (E. Evans). We thank Professor David Rentz and You Ning Su for offering resources through which we were able to access repositories of curated reference recordings with reliable metadata.

## Author contributions

CRediT: **Yucheng (Scarlett) Zhu:** Conceptualization, Data curation, Formal analysis, Investigation, Methodology, Validation, Visualization, Writing – original draft, Writing – review & editing; **Lin Schwarzkopf:** Conceptualization, Methodology, Project administration; Supervision; Writing – review & editing; **Myles H. M. Menz:** Conceptualization, Supervision; Writing – review & editing; **Slade Allen-Ankins:** Conceptualization, Data curation, Formal analysis, Investigation, Methodology, Validation, Visualization, Writing – review & editing.

## References

Alexander RD. 1960. Sound communication in Orthoptera and Cicadidae. Animal sounds and communication. 7:38–92.

Allen-Ankins S, Hoefer S, Bartholomew J, Brodie S, Schwarzkopf L. 2025. The use of BirdNET embeddings as a fast solution to find novel sound classes in audio recordings. Frontiers in Ecology and Evolution. 12:1409407.

Allen-Ankins S, Roe P, Schwarzkopf L. 2026. Needle in a haystack: Searching for threatened species using an acoustic observatory. Ecological Informatics.103850.

Bartholomew J, Schwarzkopf L, Allen-Ankins S. 2025. Using passive acoustic monitoring to investigate the occurrence of invasive Asian house geckos (Hemidactylus frenatus) on an Oceanic Island [preprint]. Research Square.

Belbin L, Wallis E, Hobern D, Zerger A. 2021. The Atlas of Living Australia: History, current state and future directions. Biodiversity Data Journal. 9:e65023.

Brodie S, Schwarzkopf L, Allen-Ankins S. 2023. Final Report to Gulf Savannah NRM Biodiversity Project [unpublished report]. James Cook University.

Digby A, Towsey M, Bell BD, Teal PD. 2013. A practical comparison of manual and autonomous methods for acoustic monitoring. Methods in Ecology and Evolution. 4(7):675–683.

Doohan B, Hoefer S, Allen-Ankins S, Nilsen V, Schwarzkopf L. 2026. Passive acoustic monitoring predicts higher avian diversity metrics than traditional bird surveys across multiple Australian bioregions. Ecological Indicators. 182:114533.

Doolan JM, Mac Nally RC. 1981. Spatial dynamics and breeding ecology in the cicada Cystosoma saundersii: the interaction between distributions of resources and intraspecific behaviour. The Journal of Animal Ecology.925–940.

Enari H, Enari HS, Okuda K, Maruyama T, Okuda KN. 2019. An evaluation of the efficiency of passive acoustic monitoring in detecting deer and primates in comparison with camera traps. Ecological Indicators. 98:753–762.

Ewart A. 2001. Emergence patterns and densities of cicadas (Hemiptera: Cicadidae) near Caloundra, south-east Queensland. TheAustralian Entomologist. 28(3):69–84.

Ewart A. 2005. Cicadas of the Pennefather River–Weipa areas, October/November 2002, with comparative notes on the cicadas from Heathlands, Cape York Peninsula. Gulf of Carpentaria Scientific Report.169–179.

Fonseca P, Allen Revez M. 2002. Temperature dependence of cicada songs (Homoptera, Cicadoidea). Journal of comparative Physiology A. 187(12):971–976.

Hart PJ, Hall R, Ray W, Beck A, Zook J. 2015. Cicadas impact bird communication in a noisy tropical rainforest. Behavioral Ecology. 26(3):839–842.

Kahl S, Wood CM, Eibl M, Klinck H. 2021. BirdNET: A deep learning solution for avian diversity monitoring. Ecological Informatics. 61:101236.

Knight EC, Hannah KC, Foley GJ, Scott CD, Brigham RM, Bayne E. 2017. Recommendations for acoustic recognizer performance assessment with application to five common automated signal recognition programs. ACE. 12(2):14.

Landis DA, Gardiner MM, van der Werf W, Swinton SM. 2008. Increasing corn for biofuel production reduces biocontrol services in agricultural landscapes. Proceedings of the National Academy of Sciences. 105(51):20552–20557.

Lavelle P, Decaëns T, Aubert M, Barot Sb, Blouin M, Bureau F, Margerie P, Mora P, Rossi J-P. 2006. Soil invertebrates and ecosystem services. European journal of soil biology. 42:S3–S15.

Leung FKW, Allen-Ankins S, Schwarzkopf L. 2026. Acoustic monitoring of invasive species: Continental patterns in calling activity of the invasive cane toads. NeoBiota. 108:89–119.

Leung FKW, Schwarzkopf L, Allen-Ankins S. 2025. Advancing invasive species monitoring: A free tool for detecting invasive cane toads using continental-scale data. Ecological Informatics. 89:103172.

Mankin RW, Hagstrum DW, Smith MT, Roda A, Kairo MT. 2011. Perspective and promise: a century of insect acoustic detection and monitoring. American entomologist. 57(1):30–44.

McGinn K, Kahl S, Peery MZ, Klinck H, Wood CM. 2023. Feature embeddings from the BirdNET algorithm provide insights into avian ecology. Ecological Informatics. 74:101995.

Mori H, Osawa T, Nitta K, Sekikawa S. 2026. Using Autonomous Recording Units to Detect Variation in Cicada Calling Pattern Across Urbanized and Green Spaces. Ecological Research. 41(3):e70066.

Moulds M. 1985. ‘Illyria’, a new genus for Australian cicadas currently placed in ‘Cicada’ L.(= ‘Tettigia’ Amyot)(Homoptera: Cicadidae). General and Applied Entomology: The Journal of the Entomological Society of New South Wales. 17:25–35.

Okamoto R, Oguma H. 2025. A simulation based approach for enhancing automatic detection of cicada songs in challenging chorus conditions. Ecological Informatics. 90:103299.

Popple LW. 2026. A web guide to the cicadas of Australia. Popple Creative Industries. https://www.dr-pop.net/cicadas.htm.

R Core Team. 2021. R: A language and environment for statistical computing [software]. Version Version 4.1.2. R Foundation for Statistical Computing.https://www.R-project.org/

R Core Team. 2025. R: A language and environment for statistical computing [software]. Version Version 4.5.0. R Foundation for Statistical Computing.https://www.R-project.org/

Rodenhouse NL, Bohlen PJ, Barrett GW. 1997. Effects of woodland shape on the spatial distribution and density of 17-year periodical cicadas (Homoptera: Cicadidae). American Midland Naturalist.124–135.

Saito T, Rehmsmeier M. 2017. Precrec: fast and accurate precision–recall and ROC curve calculations in R. Bioinformatics. 33(1):145–147.

Sanborn AF, Heath MS, Heath JE, Noriega FG. 1995. Diurnal activity, temperature responses and endothermy in three South American cicadas (Homoptera: Cicadidae: Dorisiana bonaerensis, Quesada gigas and Fidicina mannifera). Journal of Thermal Biology. 20(6):451–460.

Schowalter TD. 2016. Insect ecology: an ecosystem approach. Elsevier.

Stachowicz JJ, Bruno JF, Duffy JE. 2007. Understanding the effects of marine biodiversity on communities and ecosystems. Annu Rev Ecol Evol Syst. 38(1):739–766.

Stowell D. 2022. Computational bioacoustics with deep learning: a review and roadmap. PeerJ. 10:e13152.

Strang CA. 2025. Geography of Periodical Cicadas (Hemiptera: Cicadidae) in Portions of Northeast Illinois and Northwest Indiana. The Great Lakes Entomologist. 58(2):5.

Sueur J. 2002. Cicada acoustic communication: potential sound partitioning in a multispecies community from Mexico (Hemiptera: Cicadomorpha: Cicadidae). Biological Journal of the Linnean Society. 75(3):379–394.

Takakura K, Yamazaki K. 2007. Cover dependence of predation avoidance alters the effect of habitat fragmentation on two cicadas (Hemiptera: Cicadidae). Annals of the Entomological Society of America. 100(5):729–735.

Tolkova I, Klinck H, Cusano DA, Kügler A, Parks SE. 2025. Detection, communication, and individual identification with deep audio embeddings: A case study with North Atlantic right whales. bioRxiv.2025.2007. 2011.664307.

Traill LW, Lim ML, Sodhi NS, Bradshaw CJ. 2010. Mechanisms driving change: altered species interactions and ecosystem function through global warming. Journal of Animal Ecology. 79(5):937–947.

Wagner DL. 2020. Insect Declines in the Anthropocene. Annual Review of Entomology. 65(Volume 65, 2020):457–480.

Washnik N, Suresh C, Lee C-Y. 2023. Using Audacity software to enhance teaching and learning of hearing science course: A tutorial. Teaching and Learning in Communication Sciences & Disorders. 7(3):4.

Wolda H. 1989. Seasonal cues in tropical organisms. Rainfall? Not necessarily! Oecologia. 80(4):437–442.

Wood CM, Kahl S. 2024. Guidelines for appropriate use of BirdNET scores and other detector outputs. Journal of Ornithology. 165(3):777–782.

Wood SN. 2011. Fast stable restricted maximum likelihood and marginal likelihood estimation of semiparametric generalized linear models. Journal of the Royal Statistical Society Series B: Statistical Methodology. 73(1):3–36.

Worm B, Duffy JE. 2003. Biodiversity, productivity and stability in real food webs. Trends in Ecology & Evolution. 18(12):628–632.

Young AM. 1972. Cicada ecology in a Costa Rican tropical rain forest. Biotropica.152–159.

Young AM. 1981. Notes on seasonality and habitat associations of tropical cicadas (Homoptera:Cicadidae) in premontane and montane tropical moist forests in Costa Rica. Journal of the New York Entomological Society.123–142.

